# Underappreciated role of environmental enrichment in alleviating depression and anxiety: Quantitative evidence synthesis of rodent models

**DOI:** 10.1101/2025.06.02.657339

**Authors:** Yefeng Yang, Manman Liu, Kyle Morrison, Malgorzata Lagisz, Shinichi Nakagawa

**Affiliations:** School of Biological, Earth and Environmental Sciences, University of New South Wales, Sydney, NSW 2052, Australia; Department of Biological Sciences, University of Alberta, CW 405, Biological Sciences Building, Edmonton, AB T6G 2E9, Canada

## Abstract

Environmental enrichment has long been recognized as a non-pharmacological intervention to mitigate mental health issues, yet its efficacy, and heterogeneity of treatment effects across experimental contexts remain underexplored. Heterogeneity of treatment effects, which reflects variability in individual responses to interventions, is a critical factor in determining the generalizability and personalization needs of treatments. Here, we conducted a registered meta-analysis of 62 studies and 1,112 comparisons in rodent models to evaluate the impact of environmental enrichment on depressive and anxiety-like behaviours. We found that environmental enrichment reduced these behaviours of animal models by 16% on average and decreased inter-individual variability by 12%, indicating not only effectiveness but also low heterogeneity of treatment effects, which suggests consistent effects across individuals. Environmental enrichment further nullified the adverse effects of stressors, demonstrating a significant antagonistic interaction. These effects were robust across multiple sensitivity analyses, including model-based predictions, post-stratification, multi-model inference, publication bias correction, and critical appraisal of study quality. Moderator analyses highlighted the importance of exposure timing and the inclusion of social enrichment components. Taken together, our pre-clinical evidence on rodent models supports environmental enrichment as a low-cost, scalable, and biologically grounded intervention with translational relevance for developing equitable and accessible treatments for depression and anxiety. Given the importance of innovation and personalization in mental health care, the low heterogeneity of treatment effects of environmental enrichment positions it as a promising avenue for non-pharmacological therapeutic strategies that can be broadly applied without extensive tailoring.

## Introduction

In today’s fast-paced, often isolating world—particularly in the aftermath of the social restrictions imposed by the COVID-19 pandemic—mental health has emerged as an important public health challenge ^1–4^. Among the most prevalent and debilitating mental disorders are depression and anxiety, which threaten global health and well-being across all socioeconomic strata ^5–7^. Major depressive disorder (MDD) affects approximately 6% of the global adult population ^8^ annually and stands as the second leading contributor to chronic disease burden as measured by years lived with disability ^9^. Globally, over 322 million individuals live with depression, accounting for 7.5% of all lived with disability ^10^. Similarly, anxiety disorders constitute the largest class of mental illnesses in many Western nations ^7^, affecting over 60 million Europeans and incurring costs exceeding €74 billion per year ^11,12^.

Despite decades of progress, a substantial gap persists in the treatment approaches for mental disorders. Many individuals have little access to expensive pharmacological interventions or some have access to none at all ^13,14^. While psychotropic medications and psychotherapies remain cornerstone treatments, they have limitations. For example, antidepressants and anxiolytics often have delayed onset of action, side effects, or insufficient efficacy across symptoms^15,16^. Psychotherapies, though promising, require trained professionals and extended time commitments, making them inaccessible to many, particularly in low-and middle-income population ^17^. These realities underscore the urgent need for innovative, scalable, and equitable treatment strategies that not only alleviate mental health symptoms but are also accessible to populations historically underserved by conventional treatment systems.

Preclinical research offers a vital pathway for identifying such innovative interventions ^18,19^. Among emerging approaches, enriched environment or environmental enrichment, a non-pharmacological, cost-effective strategy involving enriched housing conditions for laboratory animals, has gained attention as a potential therapeutic tool for mental disorders in preclinical research ^15^. In rodent models, environmental enrichment typically includes enhanced sensory, social, and physical stimulation compared to standard housing, mimicking aspects of more naturalistic environments ^20^. As a behavioural and contextual intervention ^21^, an enriched environment offers the intriguing possibility of being both effective and widely implementable, especially in settings where resources for conventional treatments are limited.

Critically, an effective treatment should not only produce robust symptom reduction but also demonstrate consistency in individual-level responses. In contrast, many existing treatments for depression and anxiety are plagued by treatment resistance. Approximately 30% of individuals with major depressive disorder do not respond adequately to antidepressants, posing a formidable clinical challenge ^22^. When large inter-individual differences in treatment response, referred to as heterogeneity of treatment effects ^12,23,24^, are observed, treatment strategies often require personalization based on complex predictors of individual response ^25,26^. However, identifying and validating such predictors demands large clinical trials, usually exceeding 500 participants per arm, which are rarely feasible in mental health research ^27^.

**Figure 1.**
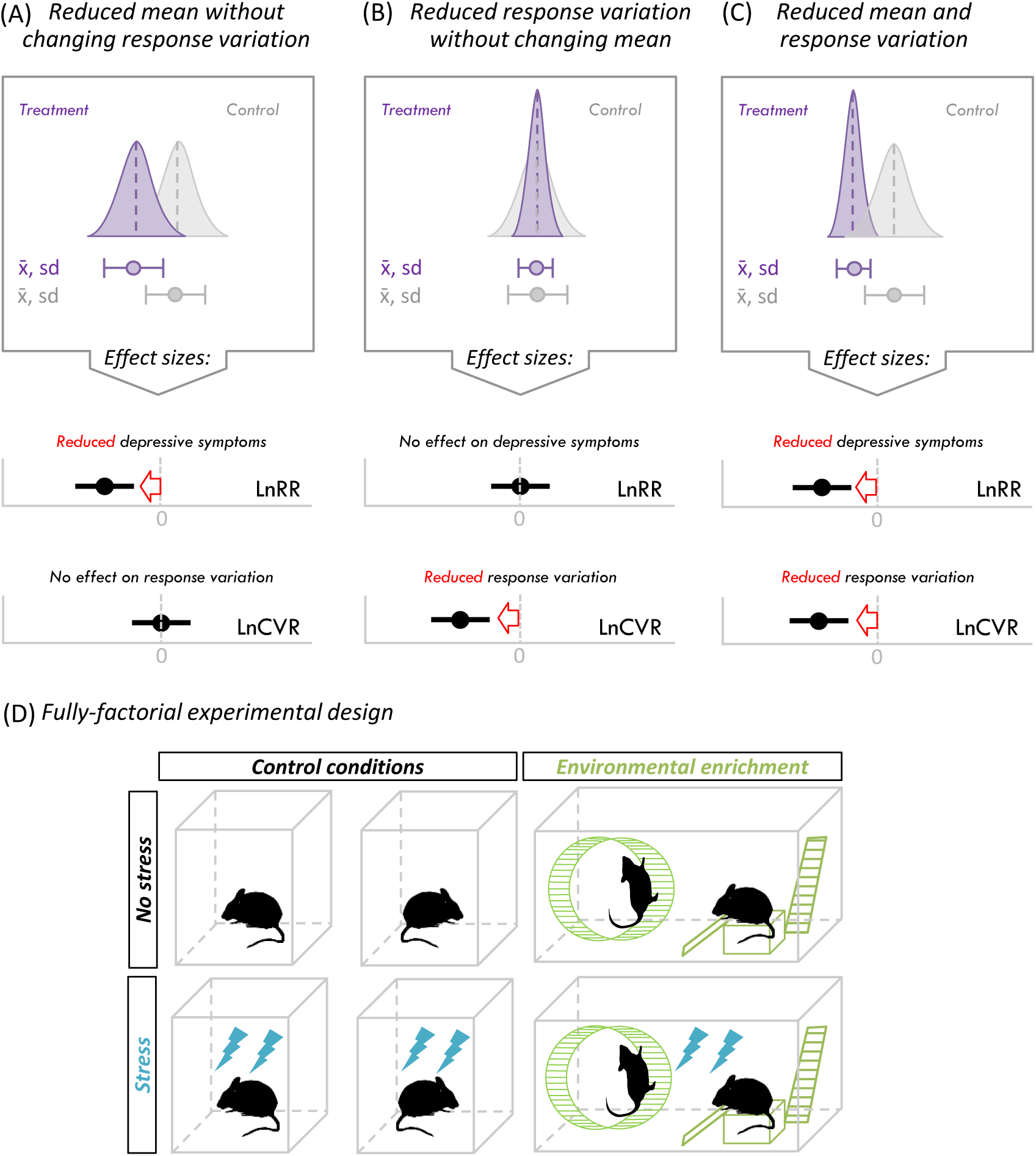
Conceptual diagram displaying three hypothetical outcomes of environmental enrichment on depressive and anxiety-like behaviours. (A) Rodents exposed to environment enrichment on average have less depressive and anxiety-like behaviours (LnRR < 0) and similar response variation (LnCVR = 0) than rodents from control group. (B) Rodents exposed to environment enrichment on average have similar depressive and anxiety-like behaviours (LnRR = 0) and less response variation (LnCVR < 0). (C) Rodents exposed to environment enrichment on average have less depressive and anxiety-like behaviours (LnRR < 0) and less response variation (LnCVR < 0). (D) Example of 2-by-2 factorial design with four groups: (1) control conditions, (2) environmental enrichment only, (3) stress only, and (4) combined environmental enrichment and stress. The log response ratio (LnRR) was used to quantify the efficacy of environmental enrichment in reducing depressive and anxiety-like behaviours (for details, see **Methods**). A negative LnRR value indicates a reduction in depressive and anxiety-like behaviours due to enrichment. The log coefficient of variation ratio (LnCVR) was used to assess the impact of enrichment on inter-individual differences (variation) in response. Negative LnCVR values suggest reduced variability or more consistent treatment effects across individuals; *x̅* is the mean values of behaviour scoresand *sd* is the standard deviation of *x̅*.

In this context, environmental enrichment presents a compelling alternative. If enriched environment consistently reduces depressive and anxiety-like symptoms across individuals, it would suggest low heterogeneity of treatment effect (Figure 1), precluding the need for highly sophisticated, individualized treatment plans ^24,28^. Recent innovations in meta-analytic methodologies allow the identification of the heterogeneity of treatment effects by examining inter-individual variability in treatment response ^28^. Specifically, the ratio of variation-based statistics between treated and control groups quantifies how interventions influence outcome variability ^29,30^. A decreased variability within the enriched environment condition would prove that the therapeutic effects do not greatly vary across individuals. Moreover, stress, a significant risk factor for mental disorders, can exacerbate depressive and anxiety-like behaviour ^31^. Whether an enriched environment retains its benefits under stress, or can even counteract stress-induced depressive and anxiety-like symptoms has not been meta-analytically tested.

Here, we conducted a pre-registered meta-analysis of 62 preclinical studies involving 12 strains of rats and mice to synthesize the effects of enriched environment on depression-and anxiety-related behaviours (https://osf.io/sg4vq/). Our findings provide the first large-scale quantitative evidence that an enriched environment not only mitigates depression and anxiety symptoms in animal models but also significantly reduces inter-individual variability in these behaviours. This evidence suggests the absence of heterogeneity of treatment effect, implying environmental enrichment may not necessitate complex personalized treatment protocols for rodents. Notably, the positive effects of environmental enrichment were observed consistently across different study designs (e.g., sex, intervention specifications), underscoring its robustness and generalizability. Furthermore, while a stressed environment predictably increased depressive and anxiety-like behaviours, the provision of an enriched environment attenuated or even nullified these effects.

## Results

### Characteristics of the evidence base

The compiled preclinical evidence base included a total of 376 effect size estimates measuring the efficacy of environmental enrichment and 281 effect size estimates assessing inter-individual differences in the response to environmental enrichment. These estimates were drawn from 116 experiments reported across 62 independent studies (see PRISMA flowchart in Figure S1). The evidence base spanned 12 of rats and mice, with Wistar rats being the most frequently used (47%), followed by Sprague-Dawley rats (18%), C57BL/6 mice (9%), and ICR mice (3%). As illustrated in Figure S2, the dataset was heterogeneous and unbalanced in its distribution of key design and population features (see below for the sensitivity analysis). A substantial majority of effect sizes (74%) were derived from male rodents. Experimental protocols varied widely in terms of the specification of enriched environments (e.g., presence or absence of social or physical enrichment components) and stress conditions (e.g., biotic vs. abiotic stressors), as well as exposure timing.

### Enriched environments reduce depressive and anxiety-like behaviours while decreasing inter-individual differences

To quantify the efficacy of environmental enrichment in reducing depressive and anxiety-like behaviours, we used the log response ratio (LnRR), calculated as the ratio of the mean behaviour score in the enriched environment group relative to the control group (for details, see **Methods**). A negative LnRR value indicates a reduction in depressive and anxiety-like behaviours due to enrichment. To assess the impact of enrichment on inter-individual differences in response, we used the log coefficient of variation ratio (LnCVR), which compares the variability (coefficient of variation; mean-standardised variation) between the enriched and control groups; negative LnCVR values suggest reduced variability or more consistent treatment effects across individuals.

**Figure 2.**
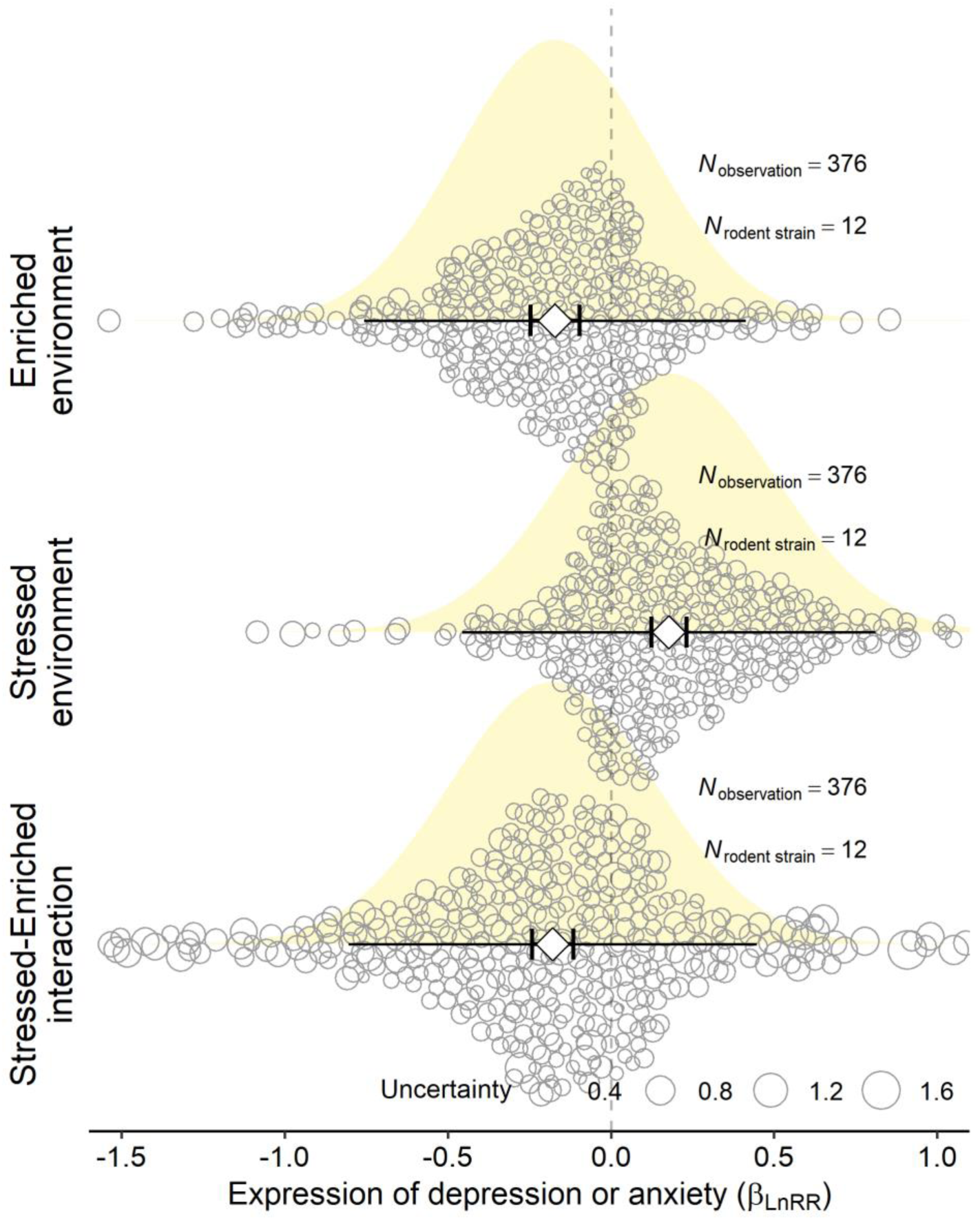
The effect of enriched, stressed, and combined environments on depression-and anxiety-like behaviours. Meta-analytic estimates (white diamonds) represent the overall mean effects (*β*_LnRR_), expressed as the log ratio of means (LnRR) between treatment and control groups, where negative values indicate reduced depression- and anxiety-like behaviours in the treatment group. Shorter whiskers denote 95% confidence intervals (CIs), while longer lines without whiskers depict 95% prediction intervals (PIs), capturing expected variation of effect sizes in future studies. Individual circles represent effect size estimates, with circle size scaled to their uncertainty (sampling variance). Shaded distributions illustrate the full prediction distributions based on the meta-analytic models ^32^. *N*_observations_ refers to the number of effect sizes included, and *N*_rodent strain_ denotes the number of distinct strains of laboratory rodent included in the meta-analytic models. For the detailed model estimate, see Table S1.

Meta-analytic results indicated that environmental enrichment significantly reduced depressive and anxiety-like behaviours by 16% (meta-analytic estimate *β*_LnRR_= -0.171, SE = 0.038, *t*(375) = -4.495, p < 0.001, 95% confidence interval, hereafter, CI [-0.246, -0.096]; Figure 2 and Table S1). Furthermore, environmental enrichment also significantly reduced inter-individual differences by 12% (meta-analytic estimate *β*_LnCVR_ = -0.127, SE = 0.051, *t*(280) = -2.494, p = 0.013, 95% CI [-0.227, -0.027]).

Despite high total heterogeneity (*I*^2^ = 96% for LnRR and 77% for LnCVR), decomposition using a multilevel meta-analytic model revealed that strain-level and between-study-level heterogeneity were both low (each <13%; Table S2), suggesting that the effects of environmental enrichment were relatively consistent across different animal strains and independent studies. The prediction distributions for strain-level and study-level effects (depicting the entire distribution of plausible effect sizes from future studies rather than merely the 95% predictive intervals ^32^) corroborated the homogeneity of the findings across these levels (Figures S3 and S4).

**Figure 3.**
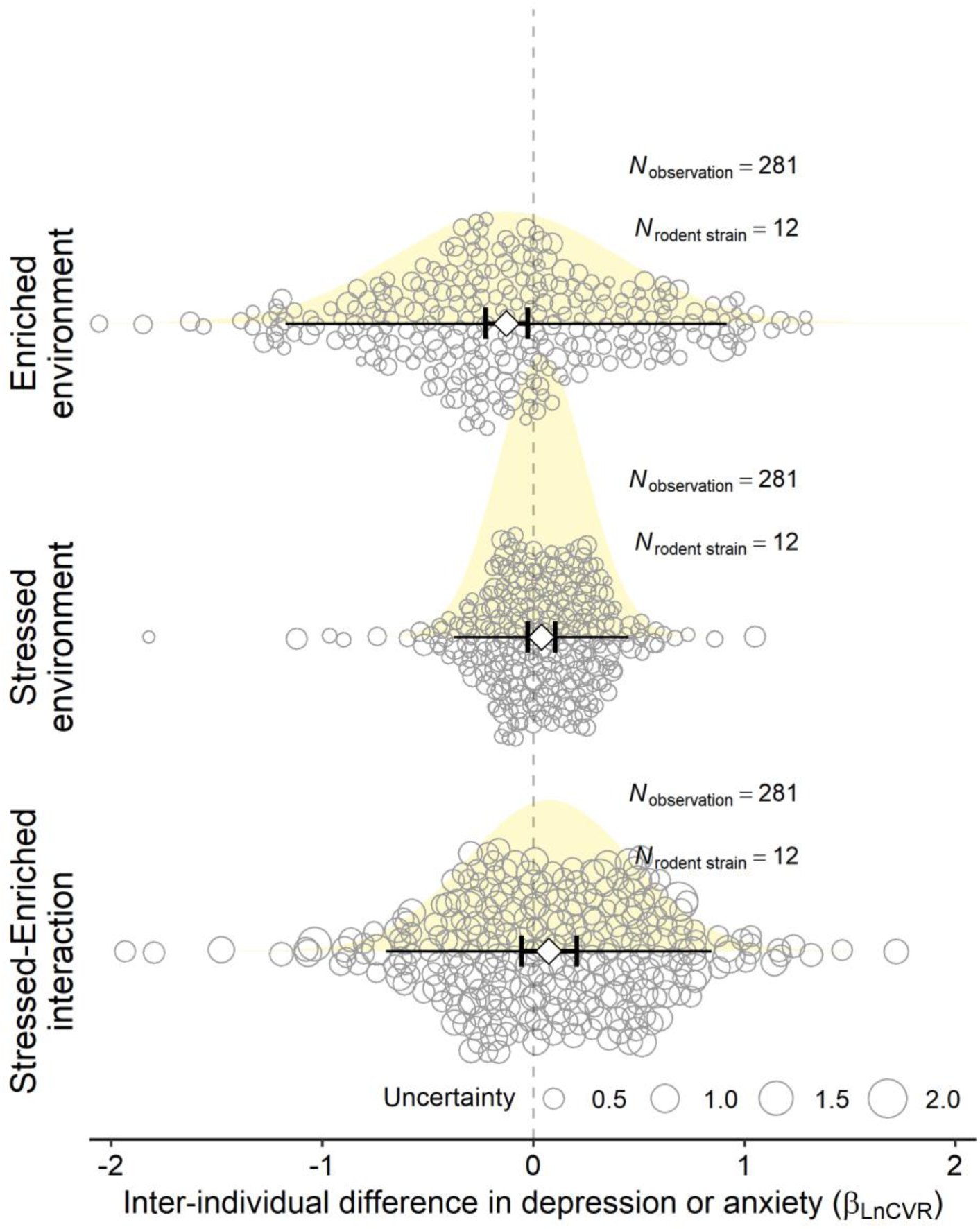
The effect of enriched, stressed, and combined environments on the inter-individual difference in depression-and anxiety-like behaviours. Meta-analytic estimates (white diamonds) represent the overall mean effects (*β*_LnCVR_) of the inter-individual variation, expressed as the log ratio of coefficient of variation (LnCVR) between treatment and control groups, where negative values indicate reduced inter-individual variation in depression-and anxiety-like behaviours in the treatment group. The remaining details are the same as in Figure 2.

### Enriched environments attenuate the effects of stressed environments on depressive and anxiety-like behaviours

We next examined whether environmental enrichment could mitigate depressive and anxiety-like behaviours in the presence of stressors (e.g., exposure to stressful environments or treatments). Exposure to stressors significantly increased depressive and anxiety-like behaviours, with an increase of 19% (*β*_LnRR_= 0.178, SE = 0.027, *t*(375) = 6.552, p < 0.001, 95% CI [0.125, 0.231]; Figure 2 and Table S1). However, when environmental enrichment was concurrently present, depressive and anxiety-like behaviours were significantly decreased by 16% (*β*_LnRR_= -0.178, SE = 0.032, *t*(375) = -5.545, p < 0.001, 95% CI [-0.241, -0.115]), indicating a strong antagonistic interaction between enrichment and stress exposure.

Importantly, the total heterogeneity for the inter-individual differences in response to combined stress and enrichment was low (*I*^2^ = 30%; Table S3), with near-zero contributions from strain-level and between-study-level heterogeneity, indicating that the antagonistic interaction effect was generalizable across different rodent strains and experimental settings. No statistically significant change in inter-individual differences was observed when enriched and stressed environments were combined (*β*_LnCVR_ = 0.075, SE = 0.067, *t*(280) = 1.126, p = 0.261, 95% CI [-0.056, 0.206]; Figure 3 and Table S1).

**Figure 4.**
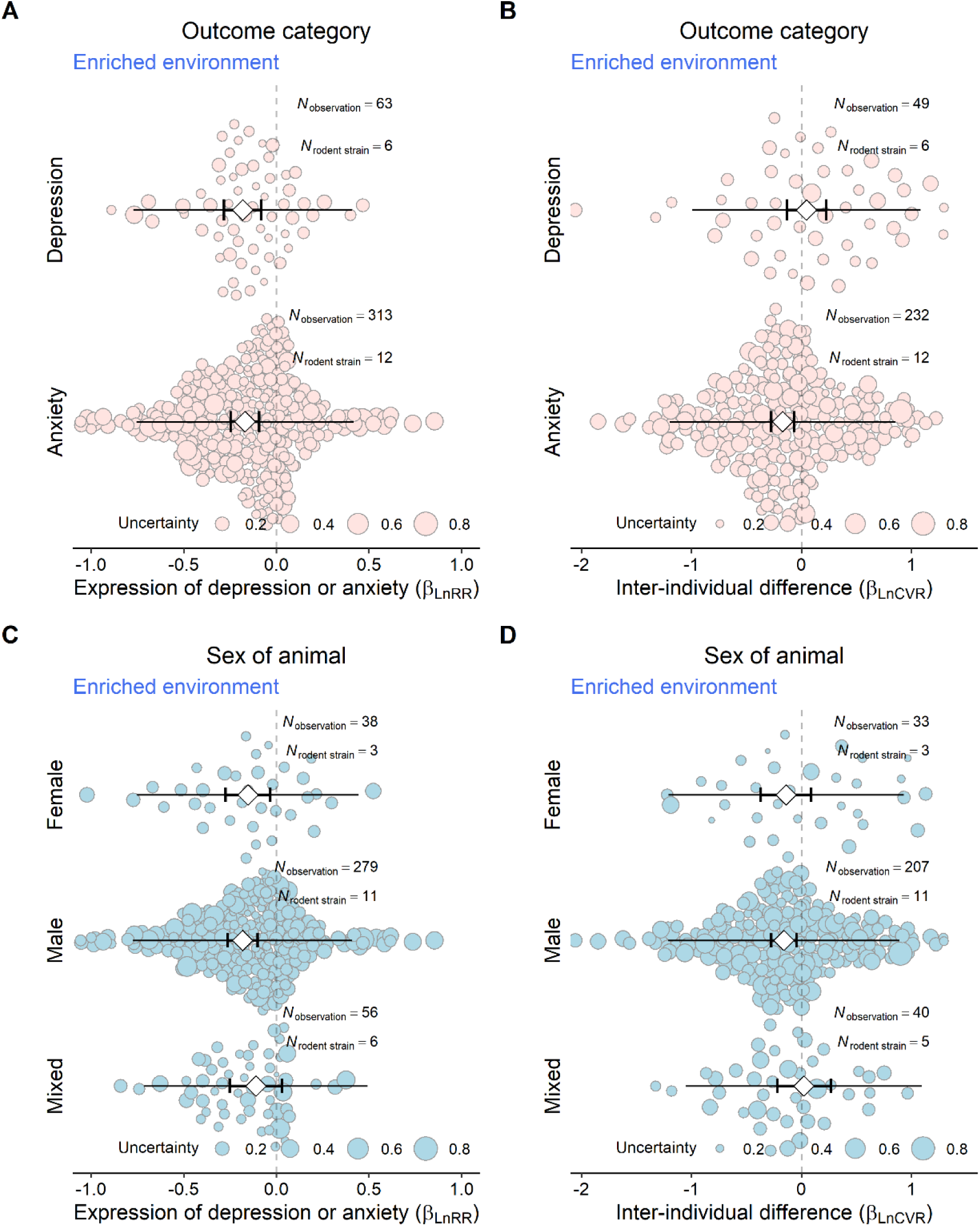
Moderator effects of behavioural outcome category and animal sex on the overall mean effects (*β*_LnRR_) and inter-individual variability (*β*_LnCVR_) of environmental enrichment. The figure illustrates how behavioural outcome categories (e.g., depression-like vs. anxiety-like behaviours) and animal sex (male, female, mixed) influence both the average treatment effect of environmental enrichment and variability in response. Effect estimates and 95% confidence and prediction intervals are shown. For the model estimates, please refer to Table S4. Further details are as described in Figure 2.

### Contextual and experimental factors drive the beneficial effects of environmental enrichment

To investigate how contextual and experimental factors influence the effects of environmental enrichment, we conducted mixed-effects meta-regression analyses on both depressive and anxiety-like behaviours (LnRR) and inter-individual variability (lnCVR). Five moderators were examined for the beneficial effect of environmental enrichment in non-stressed environments: outcome category, animal sex, exposure window of enrichment, presence of social enrichment, and presence of physical enrichment. For analyses involving the interactive effects of enriched and stressed environments, we additionally included the exposure window of stress, the presence of biotic stressors, and the presence of abiotic stressors as moderators.

All five moderators significantly influenced the beneficial effect of environmental enrichment on depressive and anxiety-like behaviours (all p < 0.001; full parameter estimates and details of statistical tests in Table S4). In the absence of stress, environmental enrichment showed a slightly stronger benefit in males than females, while mixed-sex groups did not show a significant effect (Figure 4). The timing of environmental enrichment exposure was critical: enrichment during adulthood produced the largest benefit, followed by adolescence; postnatal enrichment showed no significant effect. Social enrichment significantly enhanced the beneficial effect of environmental enrichment relative to its absence.

**Figure 5.**
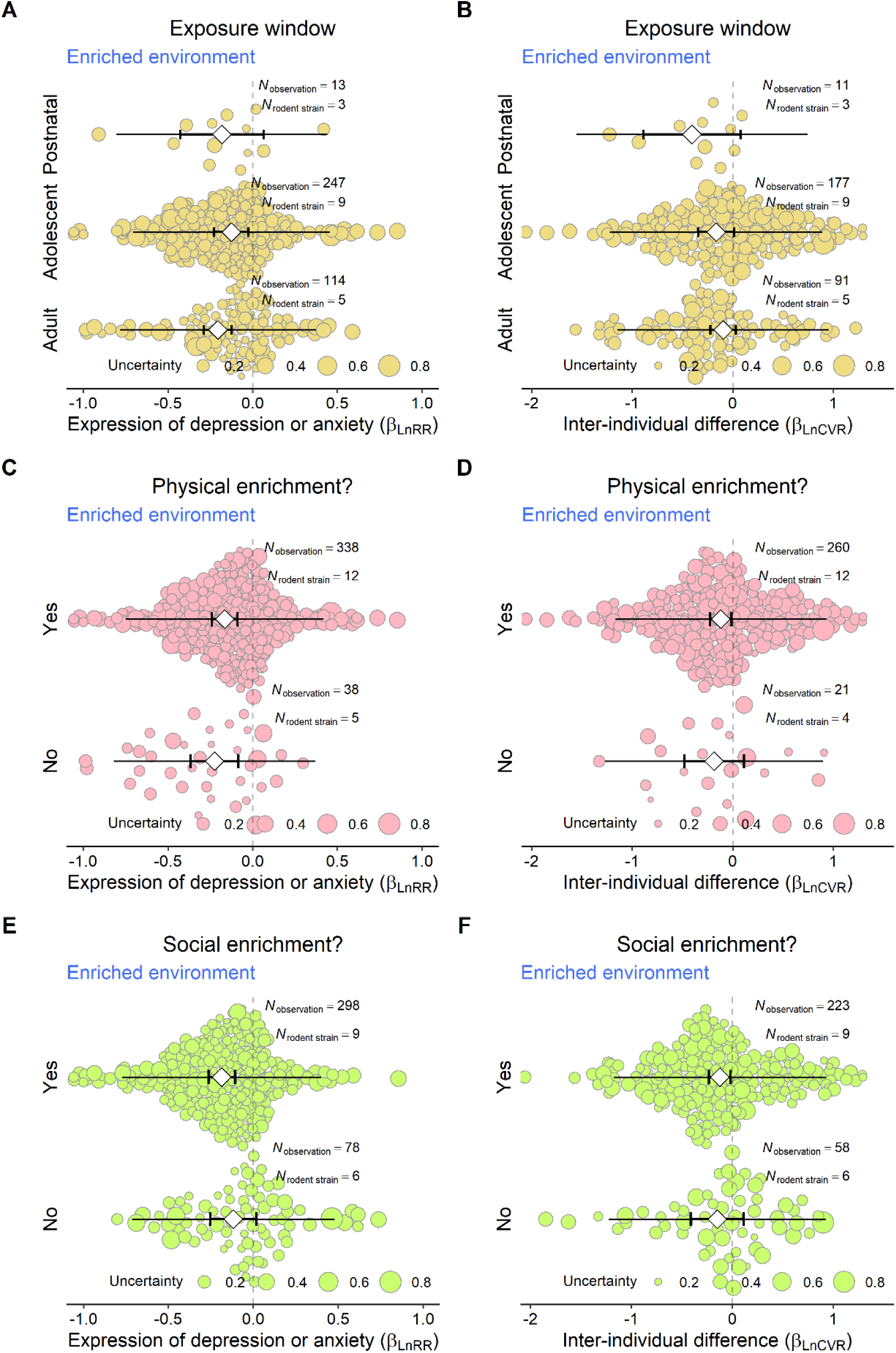
Moderator effects of enrichment- and stress-related variables on the overall mean effects (*β*_LnRR_) and inter-individual variability (*β*_LnCVR_) of environmental enrichment. The figure shows the moderating influence of the enrichment exposure window (postnatal, adolescent, adult), provision of social and physical enrichment, and presence of biotic or abiotic stressors. Results indicate how these contextual factors shape both the efficacy and consistency of enrichment effects. Effect estimates and 95% confidence and prediction intervals are displayed. For the model estimates, please refer to Table S4. Additional details are provided in Figure 2.

For inter-individual variability, all five moderators also significantly explained differences in outcomes (p-values ranging from 0.003 to 0.048). Environmental enrichment reduced variability in anxiety-like behaviours, but not in depressive-like behaviours (Figure 5 and Table S4). Variability reduction was evident in males, but not in females or mixed-sex groups. Both social and physical enrichment contributed modestly to reducing behavioural variability. In contrast, for the interaction between enriched and stressed environments, none of the eight moderators significantly influenced variability outcomes (all p > 0.25; Table S4), suggesting that the buffering effect of environmental enrichment on stress-induced depression may operate independently of these specific contextual features.

### Sensitivity analyses support the robustness of enrichment effects

To assess the robustness of the main findings, we conducted four sets of sensitivity analyses that account for potential biases related to sampling imbalance, model uncertainty, publication bias, and risk of bias. First, we examined whether the observed beneficial effects of environmental enrichment were influenced by imbalances in strain and sex representation. Our dataset was heavily skewed toward Wistar rats (47%) and male animals (74%). To mitigate the potential bias introduced by this skewed sampling, we estimated model-implied marginal mean effects under the assumption of equal representation across strain and sex categories, as well as using multilevel poststratification techniques. When assuming a balanced strain representation, the model-predicted effect size of environmental enrichment was *β*_LnRR_ = –0.148 (SE = 0.050, 95% CI [–0.244, –0.052], *t*(364) = –2.99, p = 0.003; Table S5). Assuming equal sex representation yielded an even stronger estimate (*β*_LnRR_ = –0.184, SE = 0.034, 95% CI [–0.251, –0.117], *t*(370) = –5.39, p < 0.001). Multilevel post-stratification produced similar estimates, further supporting the robustness of our conclusions regardless of sampling imbalances (detailed results in Table S5).

Second, we assessed model uncertainty via model selection and multi-model inference. Model selection using Akaike Information Criterion (AIC) indicated that several moderators contributed to improving model fit, including outcome category, sex, timing of enrichment, and the presence of physical and social enrichment. Among these, the exposure window of enrichment (e.g., adolescence versus adulthood) emerged as the most influential factor (Figure S5). Multi-model inference confirmed the significance patterns reported in the earlier moderator analysis (Table S6). The top three models, including exposure window, physical enrichment, and social enrichment, collectively accounted for 52% of the model weight among 32 candidate models.

Third, we evaluated the potential for publication bias and conducted leave-one-out diagnostics. Funnel plots adjusted for key moderators were visually symmetrical, yet Egger’s test indicated asymmetry (*t*(369) = –3.14, p = 0.002; Figure S6). After correcting for potential publication bias, the effect of enrichment on depressive and anxiety-like behaviours remained marginally significant (*β*_LnRR_ = –0.048, SE = 0.025, *t*(61) = –1.949, p = 0.056, 95% CI [–0.097, 0.001]; Figure S7). No evidence of time-lag bias was detected (*β*_LnRR_ = –0.001, SE = 0.006, *t*(369) = 0.206, p = 0.837; Figure S8), suggesting temporal stability of enrichment effects. Leave-one-out cross-validation further demonstrated that excluding any single rodent strain did not alter the estimates of the beneficial effect of environmental enrichment, confirming the robustness of the main findings (Figure S9).

Lastly, we performed a critical appraisal of methodological quality by examining subsets of studies that explicitly reported blinding or randomisation during behavioural assays. Among studies that explicitly reported blinding, enrichment significantly reduced depressive and anxiety-like behaviours (*β*_LnRR_= –0.178, SE = 0.083, *t*(81) = –2.138, p = 0.035, 95% CI [–0.343, –0.012]; Table S7). Similarly, in studies reporting randomisation, enrichment had a significant effect (*β*_LnRR_= –0.201, SE = 0.046, *t*(262) = –4.363, p < 0.001, 95% CI [–0.292, –0.110]; Table S7). These results affirm that the beneficial impact of environmental enrichment is robust even when only using evidence from low risk-of-bias studies.

## Discussion

### Environmental enrichment as a potential intervention for depressive and anxiety-like behaviours

The present meta-analysis provides robust evidence that environmental enrichment significantly reduces depressive and anxiety-like behaviours in rodent models. On average, environmental enrichment reduced these behaviours by 16%, with consistent effects across a wide range of experimental designs. This main effect underscores the potential of enrichment as a non-pharmacological intervention. Furthermore, a particularly striking finding is its antagonistic interaction with stress. Stress typically increases depressive and anxiety-like behaviours by 19%, but when environmental enrichment is introduced, it mitigates this effect, reducing the stress-induced increase by 16%. This means that enrichment does not merely alleviate symptoms in isolation; it also actively counteracts the negative impact of environmental stressors, serving as a protective buffer. This buffering capacity could be especially significant for mental health interventions, as it suggests that enrichment could enhance resilience against challenging conditions.

Primary studies, however, often struggle to consistently detect this antagonistic interaction ^33^. Their limitations stem from factors such as small sample sizes and variability in experimental conditions, which reduce statistical power and make it challenging to identify subtle or complex effects, such as the interplay between enrichment and stress. Individual experiments may yield inconsistent results due to these constraints, masking the true extent of the interaction ^34^. In contrast, the meta-analytic approach overcomes these challenges by aggregating data from 62 studies, pooling diverse findings into a larger, more representative sample. Improved statistical power allows us to detect interactive effects that may be too weak or noisy to be reliably detected by a single study ^33^. By synthesizing results across multiple experiments, the meta-analysis provides a clearer, more reliable picture of the antagonistic relationship between enrichment and stress.

The robustness of these findings is further supported by extensive sensitivity analyses, including multilevel poststratification, model selection, publication bias corrections, and critical appraisal of experimental quality. This comprehensive approach ensures that the detected interaction is not an artifact of methodological flaws or biases, reinforcing the conclusion that environmental enrichment offers consistent, generalizable benefits. For pre-clinical studies, this highlights the value of leveraging meta-analyses to uncover critical interactions that primary studies might miss, paving the way for more effective, evidence-based interventions.

### Inter-individual differences and treatment effect heterogeneity

Heterogeneity of treatment effects poses a significant challenge in mental health research, as variability in individual responses complicates the development of effective, universal interventions ^28^. Detecting heterogeneity of treatment effects in primary studies is often hindered by limited statistical power due to small sample sizes, which restricts the ability to quantify response consistency. Indeed, the statistical power primary studies to detect such variability range from 6% to 12% ^35^. Meta-analytic approaches address this limitation by pooling data across studies to enhance power and precision ^29^. For instance, studies on schizophrenia and antipsychotic response have successfully employed variance-based meta-analyses to assess heterogeneity of treatment effects ^36,37^, demonstrating the approach’s efficacy in capturing inter-individual variability ^28,38,39^.

To assess heterogeneity of treatment effects in our study, we used the natural logarithm of the coefficient of variation ratio (LnCVR), defined as *ln*(*CV_E_* / *CV_C_*), where *CV_E_* and *CV_C_* are the coefficients of variation (standard deviation/mean) for the environmental enrichment and control groups, respectively. The coefficient of variation standardizes variability relative to the mean, mitigating biases from mean-variance relationships that can distort raw variance comparisons, especially when floor effects are present ^29^. A negative LnCVR indicates lower variability in the environmental enrichment group, suggesting more consistent treatment effects or reduced heterogeneity of treatment effects (i.e., canalised inter-individual differences in response). Our finding of a 12% reduction in variability demonstrates that environmental enrichment produces more uniform outcomes compared to controls. The absence of significant moderators for LnCVR in the interactive effects of environmental enrichment and stressed environments further suggests that environmental enrichment’s stabilizing effect is robust across diverse conditions, reinforcing its potential as a generalizable intervention ^39^.

These results align with findings from McCutcheon, et al. (2021) ^40^, who investigated antipsychotic treatment response in schizophrenia. They reported that antipsychotics not only improved symptoms more than placebo (*β*_Hedge′g_ = 0.47, p < 0.001) but also reduced variability in total symptom change (*β*_LnCVR_ = -0.15, p < 0.001), contrary to their hypothesis of a non-responsive schizophrenia subtype. They interpreted a CVR < 1 (LnCVR < 0) as evidence of a more homogeneous treatment effect, suggesting that antipsychotics produce consistent benefits across patients rather than distinct responder and non-responder subgroups. Similarly, our negative LnCVR indicates that environmental enrichment yields more uniform outcomes compared to controls, challenging the notion of distinct responder subtypes in rodent models. McCutcheon, et al. (2021) ^40^, further, noted that this homogeneity reduces the need for personalized approaches, a point echoed in our study: the low heterogeneity of treatment effects, especially for anxiety, suggests that environmental enrichment could be widely applied without extensive tailoring, enhancing its translational potential as a non-pharmacological intervention.

Methodologically, McCutcheon, et al. (2021) ^40^ emphasized the importance of using CVR to adjust for mean-variance scaling, a practice we adopted to ensure reliable variability estimates. Their individual patient data analyses, which found no evidence of bimodal response distributions, further bolster our inference that environmental enrichment’s consistent effects are unlikely to mask underlying subgroups. While our preclinical context differs, the parallel use of LnCVR and the shared finding of reduced variability strengthen the interpretative framework of our results. Collectively, these insights suggest that environmental enrichment, like antipsychotics in schizophrenia, offers a relatively uniform therapeutic benefit (albeit with side effects), supporting its potential as a reliable and generalizable intervention if clinical studies confirm the benefits.

### Mechanistic insights and broader interpretation

The beneficial effects of environmental enrichment likely reflect its capacity to engage multiple neurobiological systems implicated in stress regulation, affect, and neuroplasticity ^41^. Environmental enrichment often combines physical, sensory, and social stimulation, conditions known to enhance neurogenesis, modulate hypothalamic-pituitary-adrenal (HPA) axis activity, increase brain-derived neurotrophic factor (BDNF) expression, and facilitate synaptic remodelling in key brain regions, such as the hippocampus and prefrontal cortex ^42–46^. These mechanisms offer plausible pathways through which environmental enrichment improves mood-related outcomes and resilience to stress. Importantly, our moderator analyses revealed that the exposure window of enrichment (particularly during the post-weaning stages) and the inclusion of social components are critical to its efficacy. This suggests that timing and the nature of enrichment matter, aligning with developmental neuroscience findings that critical periods of plasticity can shape long-term emotional and behavioural outcomes. Taken together, environmental enrichment may not merely compensate for environmental deprivation but actively promote adaptive neurodevelopment and mental health regulation ^47^.

### Translational relevance and policy implications

Our findings elevate environmental enrichment as a low-cost, scalable, and biologically grounded intervention with potentially significant translational relevance ^48^. In an era where mental health research increasingly emphasizes innovation, personalization, and equity, environmental enrichment emerges as a non-pharmacological approach that warrants broader attention. Compared to conventional treatments for depression and anxiety, which often involve pharmacological agents with delayed onset, side effects, and limited efficacy in treatment-resistant populations, environmental enrichment could offer an accessible and potentially preventive alternative. While preclinical, our findings suggest that enriched environments tested on laboratory rodents could inform therapeutic strategies in human settings, such as sensory, cognitive, or social stimulation programs in psychiatric care, educational institutions, or community mental health services. Environmental enrichment may also be particularly impactful in resource-constrained settings where pharmacological interventions are less feasible or acceptable. Given its capacity to reduce both symptom severity and inter-individual variability, environmental enrichment could support more equitable outcomes across diverse populations. Policymakers, funding agencies, and mental health practitioners could consider incorporating environmental enrichment principles into public health strategies, especially as part of integrative or stepped-care models for mental disorders.

### Limitations and future directions

Several limitations warrant consideration. First, while our dataset is extensive, the preclinical nature of our data necessitates caution in extrapolating to human populations, where environmental enrichment may involve more complex social and cultural factors. Translational validity, although promising, remains an empirical question requiring targeted research in human populations. Second, although we accounted for potential biases via rigorous sensitivity analyses, residual confounding due to unmeasured variables (e.g., handling procedures, housing density, experimenter effects) may still influence results. Third, our analysis focused on the presence or absence of enrichment rather than its intensity, duration, or individual components in fine-grained detail. Future studies should adopt factorial designs to disentangle the additive or synergistic effects of different enrichment elements. Lastly, behavioural outcomes were aggregated across multiple assays; while this enhances generalizability, it may obscure task-specific nuances in how EE modulates affective behaviour. Continued work integrating behavioural, neurobiological, and computational approaches will be crucial to refine enrichment-based interventions and determine their scalability in clinical and real-world contexts.

## Conclusion

Our meta-analysis demonstrates that environmental enrichment reliably reduces depressive and anxiety-like behaviours in laboratory rodents and buffers against the negative effects of stress, all while minimizing inter-individual variability. These effects are robust across experimental designs and analytical approaches, supporting the scientific validity of environmental enrichment as a behavioural intervention. Considering the growing need for innovative, accessible, and equitable treatments for mental disorders, environmental enrichment warrants greater attention as a non-pharmacological strategy with translational promise.

## Methods

We conducted a pilot test (by YY and ML) prior to the main data collection and analysis to refine our search strategy and data extraction processes, ensuring the feasibility and accuracy of our meta-analysis. The study protocol was pre-registered on the Open Science Framework (https://osf.io/sg4vq/), detailing our objectives, methods, and analysis plan to enhance transparency and reproducibility. Given the diverse rodent strains and complex data structures causing dependency in the evidence base, we employed multilevel meta-analytic models as the primary statistical technique to quantitatively synthesize findings from primary studies. These models account for nested data structures, such as multiple effect sizes within studies, and accommodate heterogeneity across rodent models and experimental designs. Our meta-analysis adheres to the Preferred Reporting Items for Systematic Reviews and Meta-Analyses (PRISMA) statement and its extension for transparent reporting, with a PRISMA checklist provided in Table S6. To facilitate computational reproducibility, all data and code used to generate our results are publicly available in a GitHub repository (https://github.com/Yefeng0920/EE_anxiety_git).

### Database development Information sources

To ensure a comprehensive literature search, we conducted systematic searches across three bibliometric databases: Web of Science (Core Collection), Scopus, and PubMed. The PRISMA flowchart summarizing the search and screening process is presented in Figure S1. To develop a finely tuned Boolean search string, we implemented a benchmarking search sensitivity procedure. We manually selected 10 benchmark articles from different sources (listed in Table S8) and defined the sensitivity of the search string as the percentage of these articles captured. The search string was optimized over 30 iterative rounds, with details of its development provided in Table S9. Additionally, we searched grey literature through EBSCOhost and BASE, which covers sources such as conference proceedings, theses, dissertations, and institutional reports, to capture relevant unpublished or non-peer-reviewed studies. The final search strategy for each academic database is outlined in Table S10, yielding 2,786 bibliometric records on 4 April 2023. After removing duplicates using string matching algorithms in the R package synthesisr ^49^ (R code available in the GitHub repository https://github.com/Yefeng0920/EE_anxiety_git) and Rayyan’s ‘Detect duplicates’ function, we obtained 1,628 unique bibliometric records. Two researchers (YY and KM) conducted systematic step-by-step screening of these records in Rayyan, as detailed in the ‘Eligibility criteria’.

### Eligibility criteria

We formulated *a priori* eligibility criteria using the population-exposure-comparator-outcome (PECO) framework to select studies for inclusion (details see below) ^50^. Eligible studies were experimental, employing a 2-by-2 factorial design with four groups: (1) environmental enrichment only, (2) stressed environment only, (3) combined environmental enrichment and stressed environment, and (4) unexposed ‘full’ controls reared in conventional/benign housing. Studies with partially crossed designs (i.e., not including all four groups within a single experiment) were excluded, as they did not allow for the calculation of interactive effect sizes. Review articles, background papers, perspectives, opinions, commentaries, and simulation studies were also ineligible.

Eligible populations were restricted to experimental studies on captive-bred, wild-type mammals, including laboratory rodents (e.g., mice, rats), monkeys, cats, sheep, and pigs. Non-mammalian vertebrates (e.g., zebrafish, zebra finch, chickens) and invertebrates (e.g., fruit flies) were excluded. Wild mammals captured from their natural habitats were ineligible due to unknown prior exposure histories. Mammals with genetic modifications or significant physical alterations (e.g., through surgery or drug injections, excluding benign manipulations like saline injections) were excluded, as such modifications could confound responses to experimental manipulations.

Studies were required to include three experimental groups within a single experiment: (1) environmental enrichment only, (2) stress only, and (3) combined environmental enrichment and stress, alongside a control group reared in conventional housing conditions (allowing benign treatments like saline injections). Eligible forms of stressors and environmental enrichment are detailed in the registered protocol (OSF link), respectively. Studies using physical alterations as stressors (e.g., teeth removal, disease induction, traumatic brain injury) or manipulating social/biotic environments without additional enrichment components (e.g., crowding, aggressive encounters) were excluded, as these could be classified as stressors. Studies involving additional exposures (e.g., chemical substances, dietary treatments, drugs) or trans-and inter-generational exposures were also excluded. Stress resulting from behavioural assays (e.g., Forced Swim Test) was not considered a stress treatment.

Main outcomes were restricted to quantitative measures of anxiety or depression behavioural phenotypes, with sufficient descriptive statistics (e.g., means, standard deviations) to calculate effect sizes for all four groups. Common assays and measures are listed in the registered protocol (https://osf.io/sg4vq/). Studies reporting only non-behavioural outcomes (e.g., gene expression, hormone levels, morphological features) were excluded.

The screening and exclusion criteria were validated during the pilot round by randomly selecting 100 studies from the retrieved records, assessed by researchers YY and ML. Detailed title, keyword, abstract, and full-text screening criteria, along with decision trees, are provided in Figure S10. After title, keyword, and abstract screening, 175 studies remained, with 35 conflicts between YY and KM resolved through discussion. Following full-text screening, 71 eligible studies were included for data extraction, with 6 conflicts resolved through discussion. The screening process is summarized in the PRISMA flowchart (Figure S1), and included studies are listed in Table S11.

### Coding procedures

We followed a registered codebook, validated using nine representative papers during a pilot stage, to standardize and facilitate data extraction from the 71 eligible studies. The extracted variables encompassed: (1) study identity information (e.g., title, publication year, DOI), (2) population attributes (e.g., species, strain, sex), (3) exposure specifications (e.g., type and timing of environmental enrichment and stress), (4) outcome characteristics (e.g., behavioural assays for depression or anxiety), (5) risk of bias information (e.g., experimental randomization, observer blinding during measurements), and (6) summary statistics (e.g., sample means, standard deviations, sample sizes). A complete list of extracted variables, including pre-defined moderator variables, is provided in registered protocol. Descriptive statistics for depressive and anxiety-like behaviours were retrieved directly from the text, tables, or supplementary data of each study. When data were presented only in figures, we digitized them using WebPlotDigitizer software (v4.6), ensuring accurate extraction of means and standard deviations. For each study, we extracted data for all four groups (environmental enrichment only, stress only, combined enrichment and stress, and control) to enable calculation of main and interactive effect sizes.

To quantify the impact of environmental enrichment on depression and anxiety, as well as inter-individual variability, we extracted five moderator variables: (1) outcome category (Depression vs. Anxiety), (2) animal sex (Male, Female, Mixed), (3) exposure window of enrichment (Postnatal, Adolescent, Adult), (4) presence of social enrichment (Yes vs. No; defined as inclusion of social stimuli such as group housing or interaction opportunities), and (5) presence of physical enrichment (Yes vs. No; defined as inclusion of physical stimuli such as running wheels or climbing structures). For the interactive effects of enriched and stressed environments, we extracted three additional moderator variables: (1) exposure window of stress (Postnatal, Adolescent, Adult), (2) presence of biotic stress (Yes vs. No, e.g., social defeat), and (3) presence of abiotic stress (Yes vs. No, e.g., restraint or unpredictable stress). These moderators were pre-defined with specific options to ensure consistency and minimize human error during extraction. Data extraction was performed by two researchers: YY extracted data from 18 papers, and MLiu extracted data from 53 papers, with all extractions by MLiu cross-checked by YY for accuracy. The piloted codebook guided the process, with moderator variables standardized using pre-defined options (e.g., for ‘presence of physical enrichment’: “yes” if a running wheel, treadmill or climbing structure was included, “no” otherwise). This approach ensured consistency across extractions and minimize human error during extraction.

### Effect size measures

To quantify the effects of environmental enrichment and stress on depressive and anxiety-like behaviours, as well as inter-individual variability, in animal models, we employed two effect size measures: the logarithm of the response ratio (LnRR) ^51^ and the logarithm of the coefficient of variation ratio (LnCVR) ^29^. These measures were calculated for two main effects (environmental enrichment and stress) and one interactive effect (between environmental enrichment and stress) across the 71 eligible studies, all using a 2-by-2 factorial design with four groups: (1) control (CT), (2) environmental enrichment only (A), (3) stress only (B), and (4) combined environmental enrichment and stress (A+B).

#### Logarithm of the response ratio (LnRR)

To quantify the impact of environmental enrichment and stress on depressive and anxiety-like behaviours, we used LnRR. We chose LnRR rather than the standardized mean difference (e.g., Hedges’ g) due to its advantages: it does not assume heteroscedasticity, often has higher statistical power, and is interpretable as the proportional change in behaviour after back-transformation ^52^. A negative LnRR indicates a reduction in depressive or anxiety-like behaviour, with the magnitude representing the percentage reduction. Our dataset included proportion data (e.g., % time spent in open arms in the Elevated Plus Maze, bounded at 0% and 100%). To address the bounded nature of such data, we applied an arcsine transformation to the means and standard deviations before calculating LnRR, ensuring normality and stabilizing variances. Parallel analyses using Hedges’ g are available in the .Rmd file in the GitHub repository (https://github.com/Yefeng0920/EE_anxiety_git).

Following Gurevitch, et al. (2000) ^53^, we derived effect size parameters for the main and interactive effects in a factorial meta-analysis. Main effects capture the net effect of one treatment across levels of the other, analogous to main effects of ANOVA ^54^. The main effect of environmental enrichment (LnRR_A_) compares the average response to enrichment (with or without stress) to the average response without enrichment. The main effect of stress (LnRR_B_) compares the average response to stress (with or without enrichment) to the average response without stress. The interaction effect (LnRR_A+B_) quantifies the extent to which the combined effect of environmental enrichment and stress deviates from the additive expectation of their individual effects, reflecting synergistic or antagonistic interactions ^54^.

For a study with population means 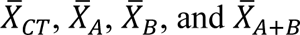 for the control, enrichment only, stress only, and combined groups, respectively, the effect size parameter for the main effect of environmental enrichment (LnRR_A_) is defined as ^55^

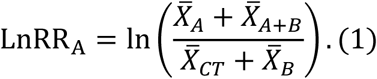

LnRR_A_ compares the average response in treatments with enrichment (A and A+B) to the average response in treatments without enrichment (CT and B). Similarly, the main effect of stress (LnRR_B_) is given by

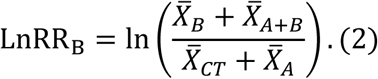

LnRR_B_ compares the average response in treatments with stress (B and A+B) to the average response in treatments without stress (CT and A). The interaction effect (LnRR_A+B_) is expressed as

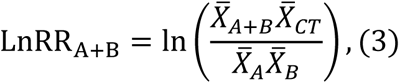

Comparing the combined effect (A+B) to the expected effect under additivity (based on A and B along relative to CT), where a non-zero value indicates an interaction between enrichment and stress (e.g., antagonistic if negative, synergistic if positive).

Estimators for these effect size parameters were obtained by substituting sample means (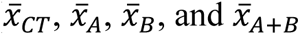; arcsine-transformed for proportion data). Sampling variances for LnRR estimators were derived using the Delta method (i.e., first-order Taylor expansion). For a log-ratio 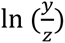, the sampling variance is approximately 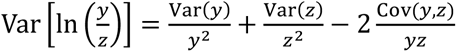. Let different treatments be mutually independent (which will be relaxed by using robust variance estimation; see ‘Meta-analytical modelling approaches’), and plugging-in sample means, sample variances 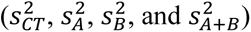 and sample sizes (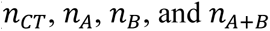) led to the estimators for sampling variance:

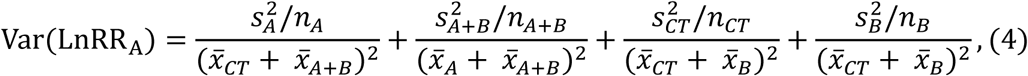

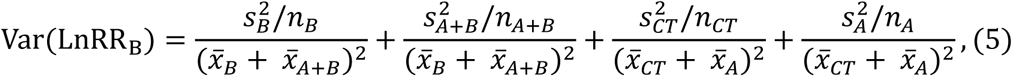

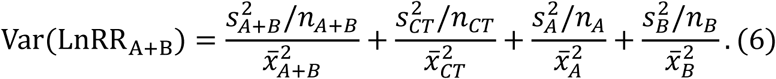

#### Logarithm of coefficient of variation ratio (LnCVR)

To quantify inter-individual variability, we used the LnCVR ^29,56^. We chose LnCVR over the logarithm of the variance ratio (lnVR) because a strong mean-variance correlation (above Pearson correlation coefficient > 0.9; Figure S11) in our dataset could confound variability comparisons through mean changes ^29^ (but see refs ^57,58^ for the limitation). By using the coefficient of variation (CV = standard deviation/mean), LnCVR standardizes variability, mitigating biases from floor or ceiling effects. Proportion data (e.g., % time in open arms) were excluded from LnCVR calculations, as CV is less meaningful for bounded data where variance is constrained near boundaries (0% or 100%). A negative LnCVR indicates reduced variability in the treatment group, suggesting lower heterogeneity of treatment effects.

The derivations of LnCVR effect size parameters followed the same factorial design principles as outlined above for the mean effects ^53^. The main effect of environmental enrichment (LnCVR_A_) compares average CV with enrichment to without enrichment, across stress levels. The main effect of stress (LnCVR_B_) compares CV with stress to without stress, across enrichment levels. The interaction effect (LnCVR_A+B_) assesses whether combined treatments alter variability beyond additivity, derived similarly to LnRR_A+B_ by comparing observed versus expected CV ratios. The effect size parameters for LnCVR are defined as:

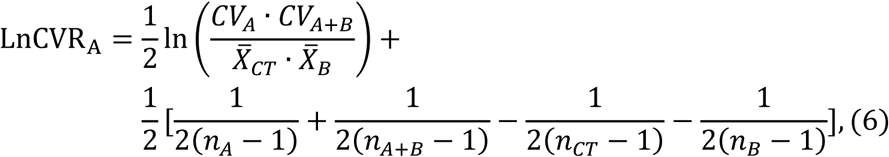

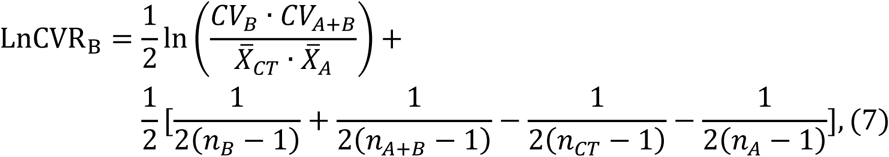

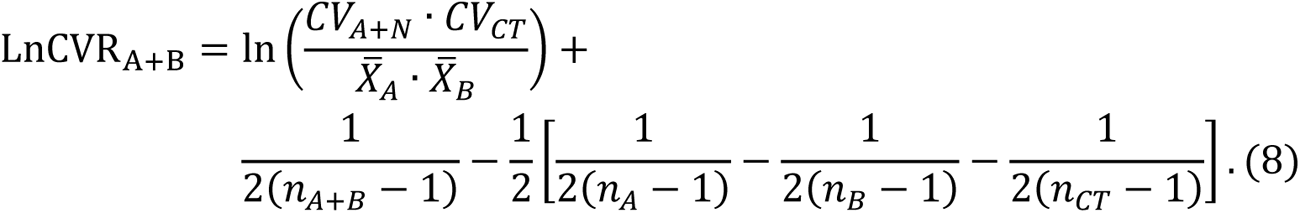

We used the sample CV (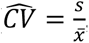, excluding proportion data) to obtain estimators. Sampling variances for the estimators were approximated by 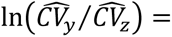 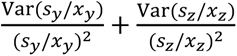. Assuming independence, and using 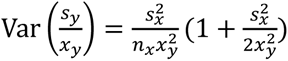, the sampling variances were derived (details see “Supplementary Methods” of Supplementary Materials).

### Statistical analysis

To quantitatively synthesize the effects of environmental enrichment and stress on depressive and anxiety-like behaviours in animal models, as quantified by LnRR and LnCVR, we employed a hybrid meta-analytic approach that accounts for the complex dependency structures inherent in our dataset. We used multilevel meta-analytic models with a sampling variance–covariance matrix, implemented within the framework of robust variance estimation (RVE) to analyse LnRR and LnCVR ^59,60^. This approach accommodates the hierarchical and correlated nature of the data, which arises from animal studies reporting multiple effect sizes. Across the 71 eligible studies, the median number of effect sizes per study was 4 (IQR = [2, 8], mean = 6), reflecting the complexity of the data. Specifically, studies often reported multiple eligible results for the same cohort (e.g., shared controls, time series measurements in one behavioural assay, or across different behavioural assays), leading to a correlated effects dependency structure. Each animal population per study contributed 4 effect size estimates, resulting in correlated sampling errors. Additionally, studies frequently reported results across non-overlapping samples (e.g., different cohorts within the same study undergoing similar assays), creating a nested effects dependency structure. Although these effect sizes are derived from independent samples, the shared study-level factors (e.g., assay protocols, measurement techniques) introduce dependency among true effect sizes within the same study.

To address these dependencies, our model specification incorporated random effects at three levels: animal-specific (to account for within-cohort correlations), study-level (to capture clustering effects), and observation-level (to model residual heterogeneity). Following the R-style symbolic expression proposed by O’Hara (2009) ^61^ for hierarchical models, the multilevel meta-analytic model used in our study can be represented as:

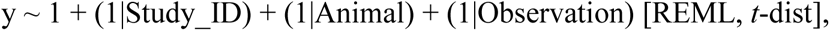

where y is the effect size (LnRR or LnCVR), 1 denotes the fixed intercept, and (1|Study_ID), (1|Animal), and (1|Observation) indicate random intercepts for study, animal strains, and observation levels, respectively. We used the restricted maximum likelihood (REML) used to estimate the variance components, and *t*-distribution (*t*-dist) with *k*−1 degrees of freedom to test the intercept (*k* is the number of observations, i.e., effect sizes).

We also included a sampling variance–covariance matrix to explicitly model the correlated sampling errors arising from shared controls or repeated measurements. The choice of this multilevel structure was partially informed by an AIC-based information-theoretic approach, which demonstrated that models with these random effects provided better fits than traditional random-effects models. To mitigate potential model misspecification, we employed RVE with a small-sample adjustment (CR1) for hypothesis testing of parameter estimates (meta-analytic estimate *β*_LnRR_and *β*_LnCVR_) and constructing confidence intervals, ensuring robust inference ^62,63^.

For moderator analysis, we used mixed-effects meta-regression by adding moderator variables (detailed in the ‘Data Items’ section) as predictors to the meta-analytic models. Given the unbalanced nature of the moderator variables, we adopted a uni-moderator meta-regression approach, including one moderator at a time to avoid overfitting. The mixed-effects meta-regression can be represented as:

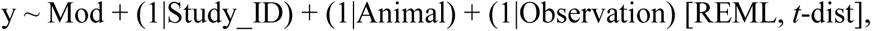

where Mod is the included moderator variable. Tests of individual parameter estimates (*β*_LnRR_and *β*_LnCVR_) were conducted using a *t*-distribution with *k*−*p* degrees of freedom, where *p* is the total number of model coefficients. Tests of the moderator effects (joint hypothesis tests of all levels of a given moderator variable) were based on an F-distribution with *m* and *k*−*p* degrees of freedom, where *m* is the number of coefficients tested. All models were fitted using the metafor package (v4.7.53) ^64^ in R (v4.0.3). All statistical tests were two-tailed and the critical value for statistical significance was *α* = 0.05. We used restricted maximum likelihood as the variance component estimator and the Quasi-Newton method to maximize the likelihood function over model parameters (embedded in metafor package). All models achieved convergence under default settings (of v4.7.53), and parameter identifiability was confirmed by visually examining the profile likelihood.

To quantify the total heterogeneity and the heterogeneity corresponding to each random effect level (animal-specific, study, and observation), we calculated *I*^2^, a metric that expresses the percentage of total variability in effect estimates due to random effects rather than sampling error ^65^. We also constructed prediction distributions to predict the range of effect sizes in new studies, following the approach of Yang, et al. (2025) ^32^, who emphasize interpreting prediction intervals and distributions to decode biological generality in meta-analyses. These prediction distributions were derived based on the total heterogeneity (in this case, variance components) and the heterogeneity at the animal and study level random effects, providing insights into the generalizability of our findings across diverse rodent populations and experimental conditions ^32^. Model specifications, including R code and data, are available in the GitHub repository (https://github.com/Yefeng0920/EE_anxiety_git) to facilitate reproducibility.

### Sensitivity analysis

To assess the robustness of our main findings on the effects of environmental enrichment and stress on depressive and anxiety-like behaviours in rodent models, we conducted four sets of sensitivity analyses, addressing potential biases related to sampling imbalance, model uncertainty, publication bias, and risk of bias. First, we examined whether the observed beneficial effects of environmental enrichment were influenced by imbalances in strain and sex representation within our dataset, which was heavily skewed toward male animals (74%) and Wistar rats (47%). To address this potential bias, we employed multilevel regression with poststratification ^66,67^, a two-step process designed to adjust effect size estimates to reflect a target population distribution. In the first step, we fitted multilevel (mixed-effects) meta-analytic regression to the LnRR effect sizes, including strain and sex as categorical moderators. This step modelled the expected effect sizes for each poststratification cell, defined by unique combinations of strain and sex levels.

We then adjusted the model-based effect size predictions to align with a target population distribution rather than the skewed empirical distribution in our dataset. For each strain (across 11 levels) and sex (across 2 levels), we computed predicted effect sizes for marginal effect estimation. We assigned poststratification weights to these predictions, reflecting equal representation across strains and sexes (50% males and 50% females), ensuring that the weights summed to one, thereby representing the target population proportions. The poststratified mean effect was calculated by aggregating these weighted predictions across all cells, using population cell sizes derived from the target distribution. To estimate the uncertainty of this poststratified mean, we approximated its variance by applying the Delta method, which leverages the covariance matrix of the predicted effects to derive a standard error for the poststratified estimate. Finally, we conducted a Wald-type hypothesis test on the poststratified mean to compute a two-tailed p-value.

Second, we assessed model uncertainty through model selection and multi-model inference ^59,68^. We fit multilevel meta-analytic models with maximum likelihood estimation, including all possible combinations of predictors (sex, exposure window, physical enrichment, social enrichment, and outcome category). This resulted in a comprehensive set of 32 candidate models. Model selection was based on the corrected Akaike Information Criterion (AICc), identifying the ‘best’ models as those with a delta AICc less than 2 compared to the top model (but see ref ^59^). To account for model uncertainty, we performed multi-model inference by averaging parameter estimates across these top models, weighted by their AICc values, and calculated the relative importance of each predictor across the entire model set.

Third, we evaluated the potential for publication bias and conducted leave-one-out diagnostics to test the influence of individual studies. For publication bias, we examined the LnRR using a contour-enhanced funnel plot (after accounting for the moderators included in the ‘best’ model) to visually assess asymmetry ^69^, which could indicate selective reporting or bias toward statistically significant results. We further tested for publication bias by including the standard errors of the LnRR as a moderator variable in a multilevel meta-regression model, analogous to the Egger regression method, to statistically detect funnel plot asymmetry ^59^. Given the Egger regression method is underpowered to detect the publication bias, we used a fixed-effect multivariate meta-analytic model with a sampling variance-covariance matrix to correct for the publication bias ^70^. Additionally, leave-one-out diagnostics involved removing each study one at a time and re-estimating the meta-analytic models to identify any single study disproportionately influencing the overall effect size estimates.

Lastly, we performed a critical appraisal of methodological quality and conducted a subgroup analysis focusing on subsets of studies that clearly reported blinding or during behavioural assays. By comparing effect size estimates between studies with and without these quality indicators, we assessed the potential impact of risk of bias on our findings, examining whether the results were driven by methodological shortcomings in the primary studies.

## Deviation

The current study had three deviations from the registered protocol: two additional analyses and one omitted analysis. These deviations were made to enhance the robustness of our findings, given unforeseen characteristics of the collected data and following best practices in meta-science. First, we introduced multilevel regression with post-stratification as an additional sensitivity analysis to address imbalances in the dataset. Second, we incorporated model selection and multi-model inference to assess model uncertainty. When we registered our protocol, we did not anticipate the imbalances in the final dataset. These imbalances, particularly in sex and strain representation, posed a risk of bias in the estimated effect sizes (LnRR and LnCVR).

To ensure the robustness of our results under a more balanced population structure, we implemented multilevel regression with poststratification to adjust for these imbalances by reweighting effect sizes based on a target population distribution. Similarly, model selection and multi-model inference were added to evaluate the uncertainty of effect size estimates across different model specifications, accounting for potential variations in moderators.

The third deviation was the omission of a planned analysis to estimate the statistical power to detect typical interactive and main effects, along with their inferential errors of effect size magnitude and sign at both the sampling and synthesis levels ^71^. In the registered protocol, this analysis was intended to quantify the inferential risk of detecting effects in our meta-analysis. However, we chose not to conduct this analysis, as post-hoc power analyses are generally discouraged, except in specific contexts such as meta-science studying science itself ^72^. Given that our study focused on synthesizing biological effects rather than meta-scientific evaluation, we prioritized alternative robustness checks, such as the sensitivity analyses described above, to test the validity of our conclusions.

## Acknowledgments

YY, ML, and SN were supported by the Australian Research Council Discovery Grant (DP210100812 & DP230101248). SN acknowledges support from a Canada Excellence Research Chair (CERC-2022-00074).

## Author contributions

YY: Conceptualization; formal analysis; investigation; methodology; software; visualization; writing – original draft; writing – review and editing. MLiu: investigation, and writing – review and editing. MK: investigation, and writing – review and editing. ML: Conceptualization; visualization; investigation, and writing – review and editing. SN: Conceptualization; investigation; methodology; writing – review and editing; funding acquisition; supervision. All authors approved the final manuscript.

## Data and materials availability

The data and scripts needed to reproduce the analyses and figures are archived at GitHub repository https://github.com/Yefeng0920/EE_anxiety_git.

